# The genome of the butternut canker pathogen, *Ophiognomonia clavigignenti-juglandacearum* shows an elevated number of genes associated with secondary metabolism and protection from host resistance responses in comparison with other members of the Diaporthales

**DOI:** 10.1101/820977

**Authors:** Guangxi Wu, Taruna A. Schuelke, Kirk Broders

## Abstract

*Ophiognomonia clavigignentijuglandacearum* (*Oc-j*) is a plant pathogenic fungus that causes canker and branch dieback diseases in the hardwood tree butternut, *Juglans cinerea*. *Oc-j* is a member of the order of Diaporthales, which includes many other plant pathogenic species, several of which also infect hardwood tree species. In this study, we sequenced the genome of *Oc-j* and achieved a high-quality assembly and delineated the phylogeny of *Oc-j* within the Diaporthales order using a genome-wide multi-gene approach. We also further examined multiple gene families that might be involved in plant pathogenicity and degradation of complex biomass, which are relevant to a pathogenic life-style in a tree host. We found that the *Oc-j* genome contains a greater number of genes in these gene families compared to other species in Diaporthales. These gene families include secreted CAZymes, kinases, cytochrome P450, efflux pumps, and secondary metabolism gene clusters. The large numbers of these genes provide *Oc-j* with an arsenal to cope with the specific ecological niche as a pathogen of the butternut tree.

## Introduction

*Ophiognomonia clavigignenti-juglandacearum* (*Oc-j*) is an Ascomycetous fungus in the family Gnomoniaceae and order Diaporthales. Like many of the other species within the Diaporthales, *Oc-j* is a canker pathogen, and is known to infect the hardwood butternut (*Juglans cinerea*) (**Figure 1**). The Diaporthales order is composed of 13 families [1], which include several plant pathogens, saprophytes, and enodphtyes [2]. Numerous tree diseases are caused by members of this order. These diseases include dogwood anthracnose (*Discula destructiva*), butternut canker (*Ophiognomonia clavigignenti*-*juglandacearum*), apple canker (*Valsa mali* & *Valsa pyri*), Eucalyptus canker (*Chrysoporthe autroafricana, C. cubensis* and *C. deuterocubensis*), and perhaps the most infamous and well-known chestnut blight (*Cryphonectira parastica*). Furthermore, several species of the Diaporthales also cause important disease of crops including soybean canker (*Diaporthe aspalathi*), soybean seed decay (*Diaporthe langicolla*) and sunflower stem canker (*Diaporthe helianti*). In addition to pathogens, there are also a multitude of species that have a saprotrophic or endophytic life strategy [3]. However, the saprophytic and endophytic species have not been studied as extensively as the pathogenic species in the Diaporthales.

**Figure 1.**
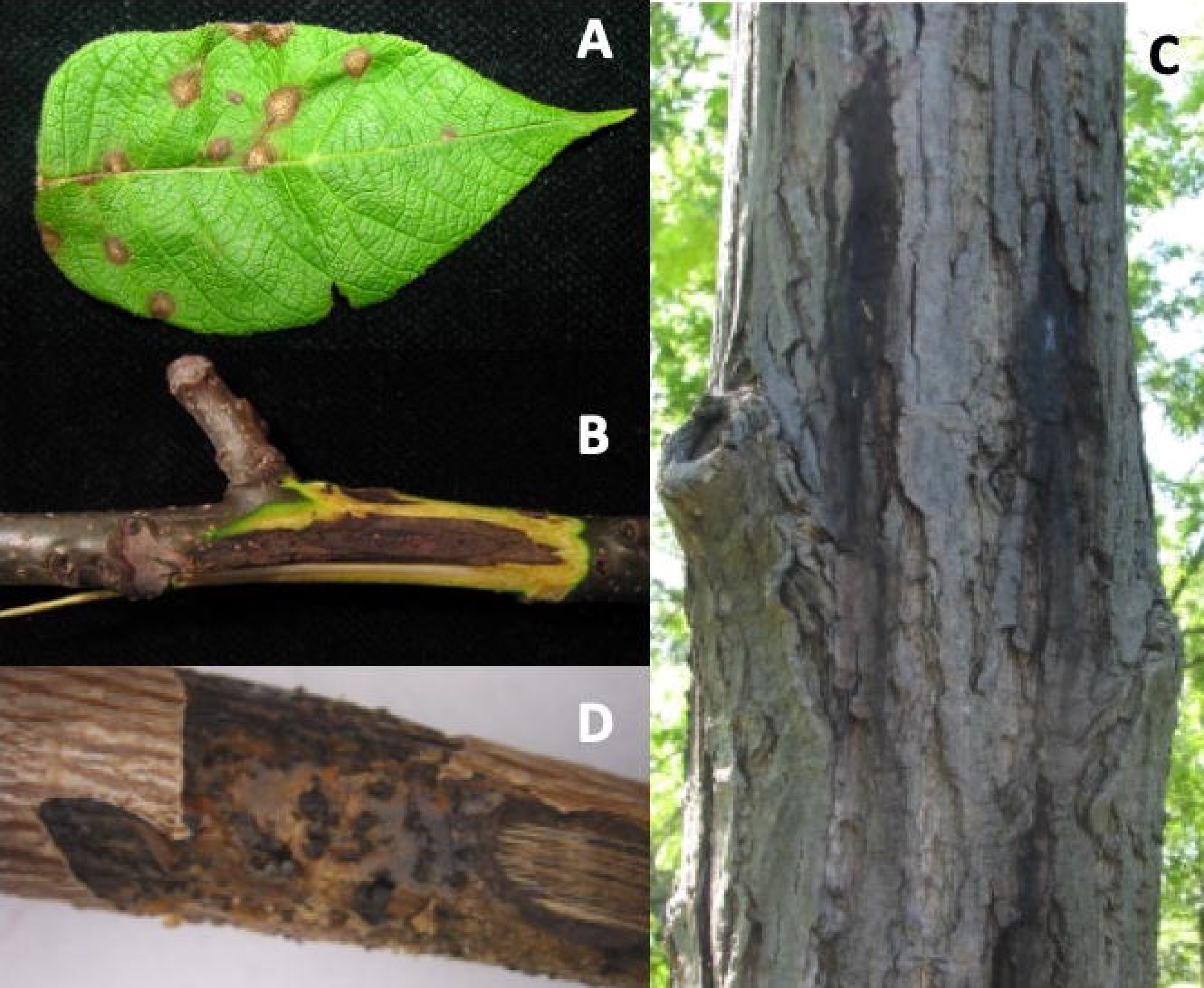
Symptoms of infection caused by Oc-j on the A) leaves, B) branches and C) trunk as well as as D) signs of fruting bodies on an infected branch.

Like many of the tree pathogens in Diaporthales, *Oc-j* is an invasive species, introduced into the U.S. from an unknown origin. This introduction caused extensive damage among the butternut population in North America during the latter half of the 20^th^ century. The first report of butternut canker was in Wisconsin in 1967 [4], and in 1979, the fungus was described for the first time as *Sirococcus clavigignenti-juglandacearum* (*Sc-j*) [5]. Recent phylogenetic studies have determined the pathogen which causes butternut canker is a member of the genus *Ophiognomonia* and was reclassified as *Ophiognomonia clavigignenti-juglandacearum* (*Oc-j*) [6]. The sudden emergence of *Oc-j*, its rapid spread in native North American butternuts, the scarcity of resistant trees, and low genetic variability in the fungus point to a recent introduction of a new pathogenic fungus that is causing a pandemic throughout North America [7].

While *Oc-j* is a devastating pathogen of a hardwood tree species, many of the species in the genus *Ophiognomonia* are endophytes or saprophytes of tree species in the order *Fagales* and more specifically the *Juglandaceae* or walnut family [8,9]. This relationship may support the hypothesis of a host jump, where the fungus may have previously been living as an endophyte or saprophyte before coming into contact with butternut. In fact, a recent study from China reported *Sirococcus (Ophiognomonia) clavigignenti-juglandacearum* as an endophyte of *Acer truncatum*, which is a maple species native to northern China [10]. The identification of the endophyte strain was made based on sequence similarity of the ITS region of the rDNA. A recent morphological and phylogenetic analysis of this isolate determined that while it is not *Oc-j*, this isolate is indeed more closely related to *Oc-j* than any other previously reported fungal species [11]. The endophyte isolate also did not produce conidia in culture in comparison to *Oc-j* which produces abundant conidia in culture. It is more likely that these organisms share are common ancestor and represent distinct species.

While the impact of members in Diaporthales on both agricultural and forested ecosystems is significant [2], there has been limited information regarding the genomic evolution of this order of fungi. Several species have recently been sequenced and the genome data made public. This includes pathogens of trees and crops as well as an endophytic and saprotrophic species [12]. However, these were generally brief genome reports and a more thorough comparative analysis of the species within the Diaporthales has yet to be completed.

Here we report the genome sequence of *Oc*-*j* and use it in comparative analyses with those of tree and crop pathogens within Diaporthales. Comparative genomics of several members of the Diaporthales order provides valuable insights into fundamental questions regarding fungal lifestyles, evolution and phylogeny, and adaptation to diverse ecological niches, especially as they relate to plant pathogenicity and degradation of complex biomass associated with tree species.

## Results and Discussion

### Genome assembly and annotation of *Oc-j*

The draft genome assembly of *Oc-j* contains a total of 52.6 Mbp and 5,401 contigs, with an N50 of 151 Kbp. The completeness of the genome assembly was assessed by identifying universal single-copy orthologs using BUSCO with lineage dataset for Sordariomycota [13]. Out of 3,725 total BUSCO groups searched, we found that 3,378 (90.7%) were complete and 264 (7.1%) were fragmented in the *Oc-j* genome, while only 83 (2.2%) were missing. This result indicates that the *Oc-j* genome is relatively complete.

### Genome-wide multi-gene phylogeny of Diaporthales

A total of 340 genes were used to generate a multigene phylogeny of the order Diaporthales. All sequenced species in Diaporthales (one genome per species for 12 species) were used. These species include: *Cryphonectria parasitica* (jgi.doe.gov), *Chrysoporthe cubensis* [14], *Chrysoporthe deuterocubensis* [14], *Chrysoporthe austroafricana* [15], *Valsa mali* [12], *Valsa pyri* [12], *Diaporthe ampelina* [16], *Diaporthe aspalathi* [17], *Diaporthe longicolla* [18], *Diaporthe helianthi* [19], and *Melanconium* sp. The outgroups used were *Neurospora crassa* [20] and *Magnaporthe grisea* [21]. We show that the order Diaporthales is divided into two main branches. One branch includes *Oc-j*, *C. parasitica,* and the *Chrysoporthe* species (**Figure 2**), in which *Oc-j* is the outlier, indicating its early divergence from the rest of the branch. The other branch includes the *Valsa* and *Diaporthe* species.

**Figure 2.**
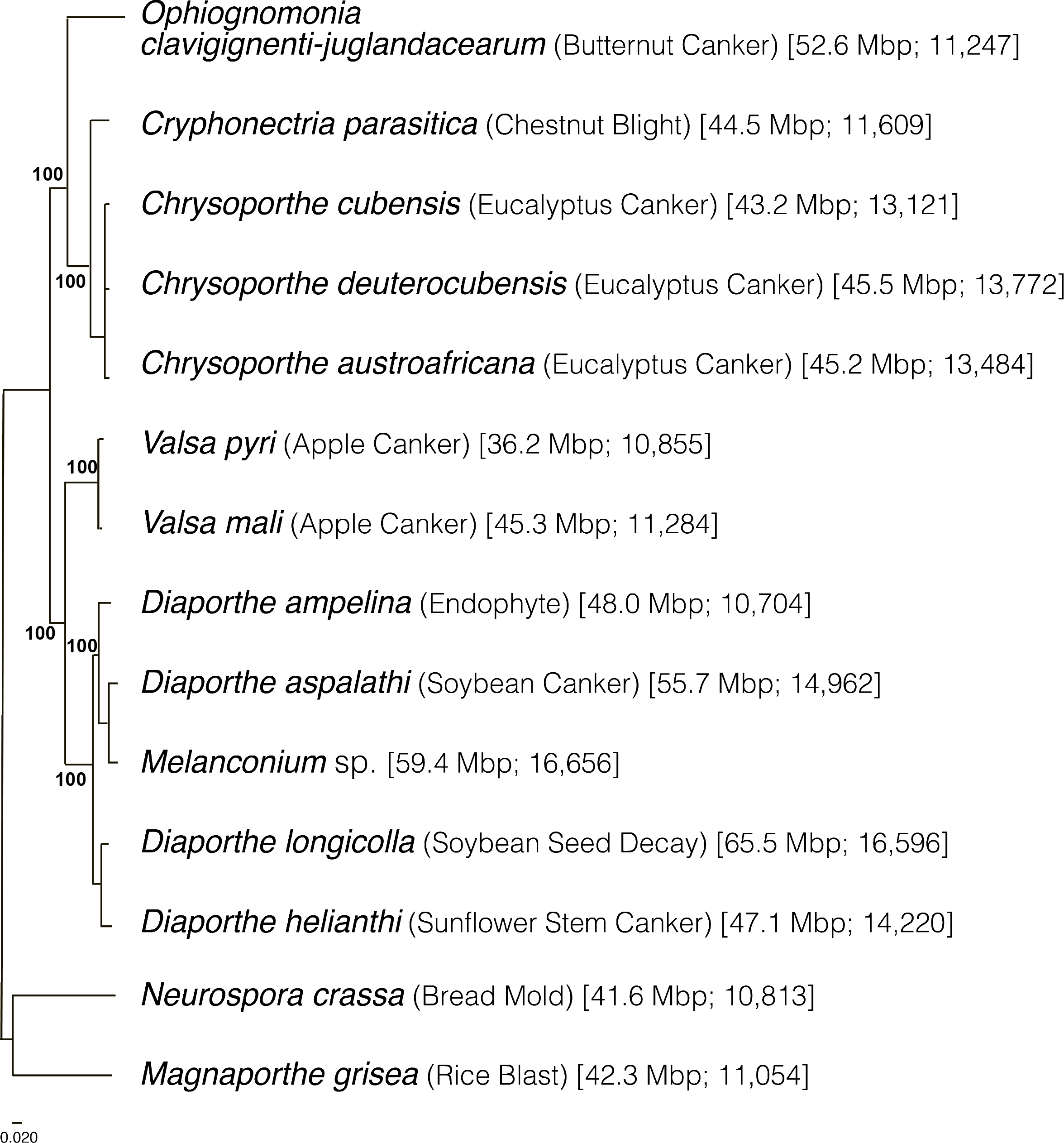
Phylogeny of Diaporthales and related species inferred using maximum likelihood by RAxML [37] with 1000 bootstraps and then midpoint rooted. The first and second numbers in parentheses represent the genome sizes in Mb and the number of predict protein models, respectively

### Gene content comparison across the Diaporthales

To examine the functional capacity of the *Oc-j* gene repertoire, the PFAM domains were identified in the protein sequences (**Supplementary Table 1**). For comparison, we also examined eight of the above mentioned 13 related species, where protein sequences were successfully retrieved (see **Methods** for details).

### Secreted CAZymes

CAZymes are a group of proteins that are involved in degrading, modifying, or creating glycosidic bonds and contain predicted catalytic and carbohydrate-binding domains [22]. When secreted, CAZymes can participate in degrading plant cell walls during colonization by fungal pathogens; therefore, a combination of CAZyme and protein secretion prediction was used to identify and classify enzymes likely involved in cell wall degradation in plant pathogens [22]. Overall, the *Oc-j* genome contains 576 putatively secreted CAZymes, more than any of the other eight species included in this analysis, and 60 CAZymes more than *D. helianthi*, the species with the second most (**Figure 3**, **Suppl. Table 1**). Given that all except bread mold *N. crassa*, which has the lowest number of secreted CAZymes, are plant pathogens, this result indicates that *Oc-j* contains an especially large gene repertoire for cell wall degradation. This pattern also holds true when compared to other tree pathogens. In a recent comparative genomic analysis of tree pathogens, it was reported that the black walnut pathogen, *Geosmithia morbida*, had 406 CAZymes, the lodgepole pine pathogen *Grosmania clavigera* had 530 CAZymes, and the plane tree canker pathogen, *Ceratocystis platani*, had 360 CAZymes [23]. Since secreted CAZymes are involved in degrading plant cell walls [22], the large number of secreted CAZymes in *Oc-j* might help facilitate its life-style infecting and colonizing butternut trees.

**Figure 3.**
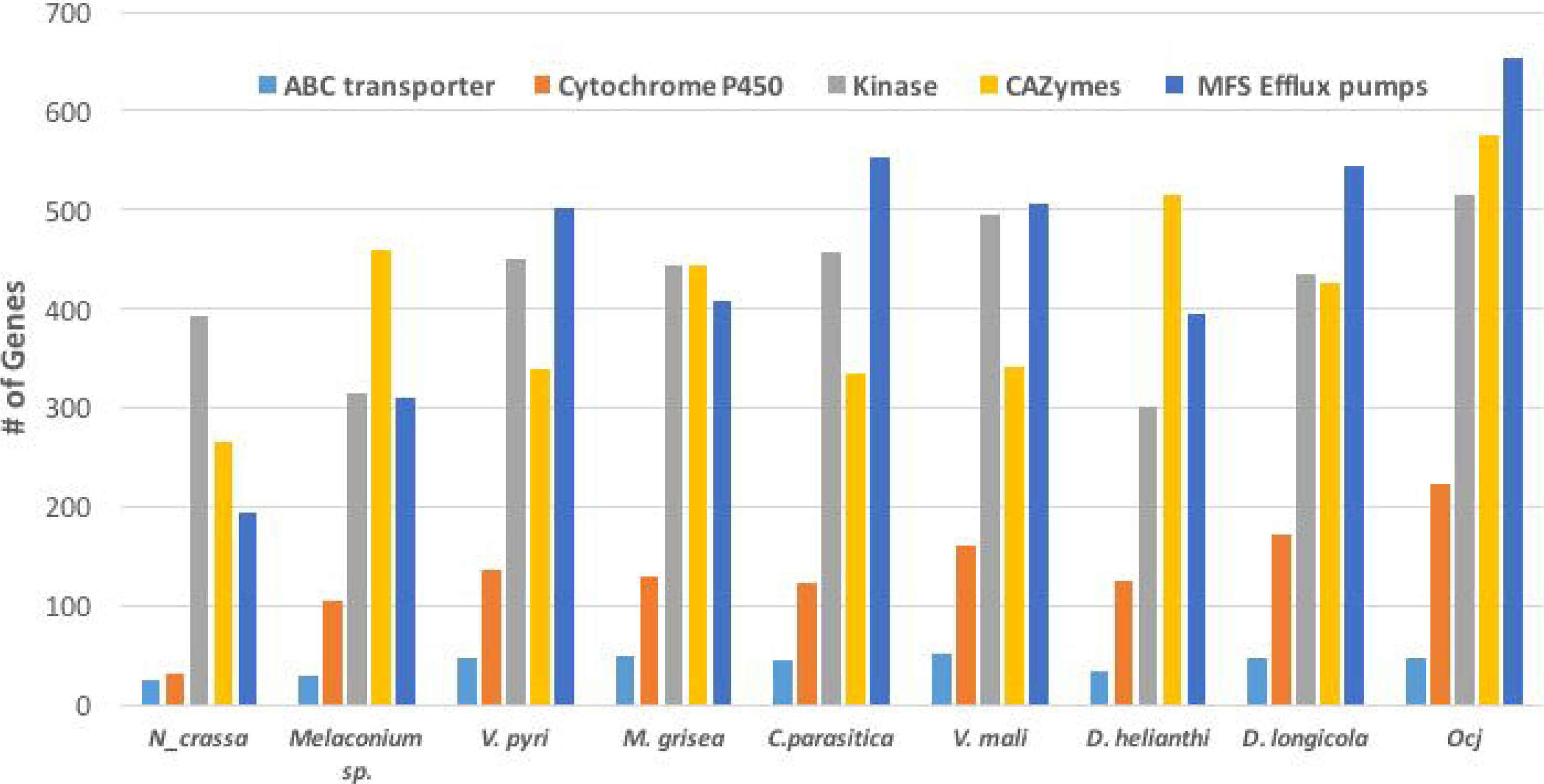
Abundance of genes in specific classes including ABC transporters, Cytochrome P450, Kinases, CAZymes, and MFS Efflux pumps present in nine species within the order Diaporthales

### Kinase

Among a total of 77 kinase related PFAM domains found in the nine above mentioned species, 61 domains are present in more genes in *Oc-j* than the average of the eight related species, while only eight domains are present in fewer genes in *Oc-j* (**Figure 3**, **Suppl. Table 1**). Given that kinases are involved in signaling networks, this result might indicate a more complicated signaling network in *Oc-j*. One example of kinase gene family expansion is the domain family of fructosamine kinase (PF03881). *Oc-j* contains 29 genes with this domain, while *N. crassa* has none and the other related species have between 3-8 genes (**Figure 3**).

### Cytochrome P450

Cytochrome P450s (CYPs) are a superfamily of monooxygenases that play a wide range of roles in metabolism and adaptation to ecological niches in fungi [24]. Among the nine species included in the PFAM analysis, *N. crassa* has only 31 CYPs, while the other eight plant-pathogenic species have between 104 – 223 CYPs. This result likely reflects that the ecological niche of *N. crassa*, bread, is more favorable for fungal growth than live plants with active defense mechanisms, thus fewer CYPs are needed for *N. crassa* to cope with relatively simple substrates. The *Oc-j* genome contains 223 CYPs (**Figure 3**, **Suppl. Table 1**), more than any of the other seven plant-pathogenic fungal species. This may be an evolutionary response to the number and diversity of secondary metabolites, such as juglone, produced by butternut as well as other *Juglans* species, that would need to be metabolized by any fungal pathogen trying to colonize the tree.

### Efflux pumps

To overcome host defenses, infect and maintain colonization of the host, fungi employ efflux pumps to counter intercellular toxin accumulation [25]. Here, we examine the presence of two major efflux pump families, ATP-binding cassette (ABC) transporters and transporters of the major facilitator superfamily (MFS) [25], in Diaporthales genomes. ABC transporters (PF00005, pfam.xfam.org) are present in all of the nine species included in the PFAM analysis, ranging from 25 genes in *N. crassa* to 51 genes in *V. mali*, while *Oc-j* has 47 genes (**Figure 3**, **Suppl. Table 1**). The major facilitator superfamily are membrane proteins which are expressed ubiquitously in all kingdoms of life for the import or export of target substrates. The MFS is a clan that contains 24 PFAM domain families (pfam.xfam.org). The *Oc-j* genome contains 653 MFS efflux pump genes, the most among all nine species included in the PFAM analysis (**Figure 3**, **Suppl. Table 1**). *C. parasitica* has 552 genes, the second most. Interestingly, it was recently shown that a secondary metabolite juglone extracted from *Juglans* spp. could be used as potential efflux pump inhibitors in *Staphylococcus aureus*, inhibiting the export of antibiotics out of the bacterial cells [26]. In line with this previous finding, our results suggest that *Oc-j* genome contains a large arsenal of efflux pumps likely due to the need to cope with hostile secondary metabolites such as the efflux pump inhibitor juglone.

### Secondary metabolism gene clusters

Secondary metabolite toxins play an important role in fungal nutrition and virulence [27]. To identify gene clusters involved in the biosynthesis of secondary metabolites in *Oc*-j and related species, we scanned their genomes for such gene clusters. We found that the *Oc-j* genome contains a remarkably large repertoire of secondary metabolism gene clusters, when compared to the closely related *C. parasitica* and the *Chrysoporthe* species (**Figure 3**, **Suppl. Table**). While *C. parasitica* has 44 gene clusters and the *Chrysoporthe* species have 18 – 48 gene clusters, the *Oc-j* genome has a total of 72 gene clusters (**Figure 3**), reflecting a greater capacity to produce various secondary metabolites. Among the 72 gene clusters in *Oc-j*, more than half are type 1 polyketide synthases (t1pks, 39 total, including hybrids), followed by non-ribosomal peptide synthetases (nrps, 14 total, including hybrids) and terpene synthases (9) (**Supplementary Table 1**).

When compared to all species included in this study, we found that *Oc-j* has the third highest number of clusters, only after *D. longicolla* and *Melanconium* sp.; however, when normalized by total gene number in each species, *Oc-j* is shown to have 6.4 secondary metabolism gene clusters per 1000 genes, the highest among all species included in this study. The *D. helianthi* genome contains only six clusters, likely due to the poor quality of the genome assembly with an N50 of ~6 kbps (**Figure 3**, **Suppl. Table 1**). Cluster numbers can vary greatly within the same genus.

## Conclusion

We constructed a high-quality genome assembly for *Oc-j*, and delineated the phylogeny of Diaporthales with a genome-wide multi-gene approach, revealing two major branches. We then examined several gene families relevant to plant pathogenicity and complex biomass degradation. We found that the *Oc-j* genome contains large numbers of genes in these gene families. These genes might be essential for *Oc-j* to cope with its niche in the hardwood butternut (*Juglans cinerea*). Future research will need to focus on understanding the prevalence of these genes associated with complex biomass degradation among other members of the *Ophiognomonia* genus which include endophytes, saprophytes and pathogens. It will be interesting to know which of these different classes of genes are important in the evolution of distinct fungal lifestyles and niche adaptation. Future research will also focus on comparing the butternut canker pathogen to canker pathogens of other tree species which produce large numbers of secondary metabolites known to be important in host defense. Our genome also serves as an essential resource for the *Oc-j* research community.

## Methods

### DNA extraction and library preparation

For this study, the type culture of *Oc-j* (ATCC 36624) recovered from an infected butternut tree in Wisconsin in 1978, was sequenced. For DNA extraction, isolates were grown on cellophane-covered potato dextrose agar for 7-10 d, and mycelia was collected and lyophilized. DNA was extracted from lyophilized mycelia using the CTAB method as outlined by the Joint Genome Institute for whole genome sequencing (Kohler A, Francis M. Genomic DNA Extraction, Accessed 12/12/2015 http://1000.fungalgenomes.org/home/wp-content/uploads/2013/02/genomicDNAProtocol-AK0511.pdf). The total DNA quantity and quality were measured using both Nanodrop and Qubit, and the sample was sent to the Hubbard Center for Genome Studies at the University of New Hampshire, Durham, New Hampshire. DNA libraries were prepared using the paired-end Illumina Truseq sample preparation kit, and were sequenced on an Illumina HiSeq 2500.

### Genome assembly and annotation

We corrected our raw reads using BLESS 0.16 [28] with the following options: -kmerlength 23 - verify ‒notrim. Once our reads were corrected, we trimmed the reads at a phred score of 2 both at the leading and trailing ends of the reads using Trimmomatic 0.32 [29]. We used a sliding window of four bases that must average a phred score of 2 and the reads must maintain a minimum length of 25 bases. Next, *de novo* assembly was built using SPAdes-3.1.1 [30] with both paired and unpaired reads and the following settings: -t 8 -m 100 --only-assembler.

Genome sequences were deposited at ncbi.nlm.nih.gov under Bioproject number PRJNA434132. Gene annotation was performed using the MAKER2 pipeline [31] in an iterative manner as is described in [32], with protein evidence from related species of *Melanconium* sp., *Cryphonectria parasitica*, *Diaporthe ampelina*, (from jgi.doe.gov) and *Diaporthe helianthi* [19], for a total of three iterations. PFAM domains were identified in all the genomes using hmmscan with trusted cutoff [33]. Only nine species were included in the PFAM analysis. For the three *Chrysoporthe* species, *D. aspalathi*, and *D. longicolla*, protein sequences were not readily available for downloading online and e-mail requests were unsuccessful. Secondary metabolism gene clusters were identified using antiSMASH 4.0 [34].

### Species phylogeny

Core eukaryotic proteins identified by CEGMA [35] were first aligned by MAFFT [36] and then concatenated. Only proteins that were present in all genomes and all sequences were longer than 90% of the *Saccharomyces cerevisiae* ortholog were used. Phylogeny was then inferred using maximum likelihood by RAxML [37] with 100 bootstraps and then midpoint rooted.

### Secreted CAZyme prediction

SignalP [38] was used to predict the presence of secretory signal peptides. CAZymes were predicted using CAZymes Analysis Toolkit [39] based on the most recent CAZY database (www.cazy.org). Proteins that both contain a signal peptide and are predicted to be a CAZyme are annotated as a secreted CAZyme.

## Acknowledgements

Special thanks to Dr. Matt MacManus. This started as an in-class project for PhD student TS and provided the basis for this final manuscript. Dr. Broders was supported by the Simon’s Foundation Grant number 429440 to the Smithsonian Tropical Research Institute.

---

**Supplementary Table 1. PFAM and antiSMASH annotation of *Oc-j* and related species**

